# Wheat cells show positional responses to invasive *Zymoseptoria tritici*

**DOI:** 10.1101/2022.07.15.499463

**Authors:** Francesco Valente, Jessica Mansfield, Daniel Herring, Giuseppe Romana, Cecilia Rodrigues, Jeremy Metz, Melanie Craze, Sarah Bowden, Andy Greenland, Julian Moger, Ken Haynes, David M. Richards, Emma Wallington, Michael J. Deeks

## Abstract

The stomatal complex of grasses consists of two guard cells and two adjacent subsidiary cells that cooperate during stomatal closure. *Zymoseptoria tritici*, the main causal agent of Septoria tritici blotch in wheat, enters the host via stomata. Here we test the hypothesis that the stomatal complex shows focused sub-cellular responses to invading *Z. tritici* hyphae.

We have combined live-cell transmission light microscopy, immunofluorescence and CRS microscopy to identify cell wall modifications triggered by hyphal invasion. Furthermore, we have used confocal fluorescence microscopy and automated quantitative image analysis to assess whether host cells respond to hyphae through spatial redistribution of organelles.

We find that subsidiary cells construct papillae that are accurately aligned with hyphal position even when hyphae are occluded by guard cells. These are distinct from those induced by powdery mildew, with callose restricted to a crust that surrounds content with a high-amplitude Raman signal in the CH-band. Peroxisome populations in subsidiary cells show distributions with modes weakly correlated with hyphal position but do not differ significantly between compatible and incompatible interactions.

Our data suggest local changes to cell wall architecture and focal accumulation of organelles in subsidiary cells could play roles in crop defence during host leaf penetration by *Z. tritici*. Molecular strategies to amplify these responses may provide novel routes for crop protection.

## 1. Introduction

The ascomycete *Zymoseptoria tritici* is the main causal agent of Septoria tritici blotch (STB), which is recognised as one of the most devastating wheat foliar diseases worldwide, causing up to 20–40% yield loss (Fones and Gurr, 2015; Kema and van Silfhout, 1997). The *Z. tritici* life cycle is characterised by a symptom-less phase of several days that follows the passage of invasive hyphae through leaf stomata. During this asymptomatic phase, the invading hyphae grow intercellularly through mesophyll tissue without greatly increasing in fungal biomass before eventually entering a necrotrophic growth phase that induces host cell death (Keon et al., 2007; O’Driscoll et al., 2015). Recently it has been suggested that some *Z. tritici* strains may be primarily epiphytic during the latent asymptomatic phase with only a small proportion of fungal growth leading to the invasion of host tissue (Fones et al., 2017; Haueisen et al., 2019). During the infection’s necrotrophic phase the phytopathogen develops asexual pycnidia sporulation structures that form within substomatal cavities (Orton et al., 2011; Steinberg, 2015). Sexual airborne ascospores are generated during the intercrop seasons on infested wheat debris and represent the primary source of inoculum (Suffert et al., 2011). Both ascospores and pycnidiospores can simultaneously be involved in the initiation of disease epidemics (Morais et al., 2015).

Recent work to clone wheat genes conferring resistance to *Z. tritici* has identified two membrane-integrated receptor kinases thought to be functional at the host cell plasma membrane. *Stb6*, encoding a cell-wall associated kinase (WAK), provides resistance to *Z. tritici* strains carrying a corresponding *AvrStb6* allele (Saintenac et al., 2018; Zhong et al., 2017), and *Stb16q* encodes a plasma membrane cysteine receptor like kinase responsible for slowing *Z. tritici* penetration and intercellular growth (Saintenac et al., 2021). Taken together, this suggests that perception of fungal activity at the host cell surface is critical for successful defence. The nature of host responses to *Z. tritici* remain uncharacterised at the sub-cellular scale. Fungi and other microbes that irritate or penetrate host cells trigger sub-cellular focal responses that include cell wall reinforcement through local deposition of callose, arabinoxylan and cellulose (Hückelhoven, 2014; Voigt, 2014). Such cell wall reinforcement acts as a barrier at the pathogen site, which may or may not prevent pathogen invasion (Voigt and Somerville, 2009). In parallel to cell wall modification, internal cell dynamics show focal responses to fungi and other phytopathogens. These include cytoskeletal rearrangement at the site of challenge (Sassmann et al., 2018), accumulation of specific organelles, and localised physiological changes (Fuchs et al., 2016).These responses can be fast with rearrangements achieved within seconds of focal stimulation (Hardham et al., 2008, 2007). They can also extend across cell borders from the site of fungal interaction (Underwood and Somerville, 2008), suggesting that focal immunity can be triggered and guided intercellularly. Quantitative live-cell imaging using automated analysis methods provides a potential means to measure and compare focal responses between stimuli and host genotypes (Fuchs et al., 2016) but phytopathology in wheat has so far rarely been explored using quantitative cell biology. Consequently, it is unknown whether wheat immune responses towards *Z. tritici* show aspects of focal defence.

In this study, we tested the hypothesis that wheat epidermal cells surrounding stomata react to non-penetrative *Z. tritici* hyphae by initiating focused immune responses. The stomatal apparatus of wheat consists of lateral subsidiary cells adjacent to the guard cells, structured either as a ring or as a pair, but distinct from other epidermal cells in morphology and perhaps unique in molecular identity (Rudall et al., 2013). Recent studies have suggested that subsidiary cells function to enhance stomatal responsiveness and greater aperture range, improving plant performance during abiotic stress including drought and heat (Raissig et al., 2017). Their role in plant immunity remains uncharacterised. Here we show that subsidiary cells have the capability to detect and focally respond to pathogenic *Z. tritici* hyphae that breach the stomatal aperture. We demonstrate the presence of papillae-like structures as an indicator of cell wall response, occurring specifically at sites neighbouring hyphal tip growth but also in mirrored locations in opposing subsidiary cells. Furthermore, we have produced transgenic wheat expressing fluorescent protein fusions to label peroxisomes and have applied novel quantitative bioimaging approaches based on four-dimensional tracking of organelles to characterise the spatiotemporal arrangement of peroxisomes relative to *Z. tritici* hyphae. We present a novel methodology to examine the role of subsidiary cells in plant immunity and to validate the presence of focal immunity in wheat epidermal cells interacting with invading phytopathogens prior to host colonisation.

## 2 Materials and methods

### 2.1 Plant material and growth conditions

Wheat plants for infection assays included cultivars Bobwhite S56, Riband and Cadenza. These were grown in a Sanyo MLR352 growth cabinet with 16 h day-length (150 µmol/m^2^/s) and 8-hour dark period at 20°C before infection at 18 days after germination (DAG). *Hordeum vulgare* L. variety ‘Golden Promise’ was grown at 150 µmol/m^2^/s with a 16-hour photo period (21 °C) and 8-hour dark period (19 °C) prior to its use for powdery mildew cultivation.

### 2.2 Fungal phytopathogen strains and infection

*Z. tritici* isolates IPO323 and K4979N2X1 (a UK field isolate provided by Syngenta, Jealott’s Hill, UK) were stored at -80 °C in 50% (v/v) glycerol and were inoculated onto Yeast Peptone Dextrose agar (YPD agar, per litre: 10 g yeast extract, 20 g peptone, 20 g dextrose and 2 g agar; lab, UK) in the presence of 0.01% ampicillin (50 µg/mL). 500 µL of sterile Milli-Q H_2_O was added to the surface of the agar to enhance humidity. Cultures were grown for 4-5 days at 18 °C. Infection assays were performed as follows: Cells were harvested from the agar plates, resuspended in sterile Milli-Q H_2_O, and filtered using a 100 µm cell strainer (Fisher, UK). For plant infection, we followed the method suggested by Keon (Keon et al., 2007). Briefly, the leaves were evenly inoculated with fungal spores standardised with a haemocytometer at a density of 5 × 10^6^ spores mL^−1^ in sterile Milli-Q H_2_O containing 0.01% (v/v) Tween-20. Infected plants were kept in growth chambers at 16 h day-length (130 µmol/m^2^/s) and 8-hour dark period at 18 °C and 85% relative humidity. The infection outcome was quantified on day 24, by measuring the percentage area of leaves occupied by distributions of pycnidia. Infected leaves were detached and incubated for 24 hours in a humid environment. Leaves were then digitally scanned at 1600 dots per inch and areas measured using Fiji.

*Blumeria graminis* f. sp. *hordei* (*Bgh*), UK isolate CC/133 (NIAB, Cambridge, UK) was cultivated on *Hordeum vulgare* L. variety ‘Golden Promise’ (8 h photoperiod, 120 µmol/m^2^/s, at 17 °C) by weekly infection of two-week-old plants (Sassmann et al., 2018). Infection of wheat to observe non-host responses in epidermal cells was performed using plants of the same age and setup described above for *Z. tritici* inoculation. Conidia were gently brushed onto wheat leaves using a soft paint brush and germinated under high humidity in an autoclave bag for a period between 24 and 40 hours in the same growth environment described above for *B. graminis* growth.

### 2.3 Wheat transformation

Wheat peroxisomes were labelled using mCherry fused to a PTS1 targeting signal consisting of a short carboxy terminus tripeptide (serine-lysine-leucine; SKL) that facilitates Pex5-mediated peroxisome import. A 653bp partial mCherry-SKL coding sequence was obtained from plasmid px-rk (Nelson et al., 2007; ABRC stock CD3-983) digested with NcoI/SnaBI and ligated into binary vector pEW145-SCV (Biogemma SAS) NcoI/SnaBI sites treated with shrimp Alkaline Phosphase (SAP), to create pEW230. The mCherry-SKL coding sequence was then completed by ligation of a 435bp px-rk NcoI fragment into pEW230 digested with NcoI and treated with SAP to create pEW235. Actin filaments were labelled using GFP-Lifeact (Smertenko et al., 2010) that was cloned using a similar strategy. A 720bp partial GFP-Lifeact coding sequence was obtained from plasmid pMDC43 digested with NcoI/EcoICRI and ligated into pEW145-SCV NcoI/SnaBI sites, treated with SAP to create pEW240 and sequenced. To create a compatible restriction site at the 5’ end of the GFP-Lifeact sequence, the 5’ region was then amplified from pMDC43 with Pfu polymerase (Stratagene) and primers MDC43-for and MDC43-rev (Supplemental Table 1). The amplicon was digested with NcoI and the 167bp fragment ligated into pEW240 digested with NcoI and treated with SAP to create pEW242 and sequenced. The final binary constructs, pEW230 and pEW242, contain an *nptII* selectable marker expressed from the Subterranean Clover Mosaic Virus Sc4 promoter with AtFAD2 intron plus the OsActin promoter for expression of the mCherry-SKL or GFP-Lifeact coding sequences *in planta* (Supplemental Figure 1). Transient transformation of wheat epidermal cells was achieved using a Helios BioRad particle delivery system with particle preparation performed according to manufacturer’s instructions. Briefly, 1.5 μm gold particles were co-coated with both pEW235 and pEW242 (0.5 μg of each plasmid and 0.5 mg of gold per cartridge). The epidermal cells of detached wheat leaves were bombarded at a helium pressure of 150 psi and incubated overnight in a dark humid environment at room temperature before imaging.

*A. tumefaciens* strain LBA4404 pSB1 (Komari et al., 1996) containing binary construct pEW235 was used for transformation of hexaploid wheat cv. Bobwhite S56 (CIMMYT, Mexico). Wheat plants were grown in controlled environment chambers at 20°C day 15°C night with a 16 hour photoperiod (approximately 400 µmol/m^2^/s). Immature embryos were inoculated *in planta* 14-20 days post-anthesis using the Seed Inoculation Method (SIM) licensed from Biogemma SAS, and subsequent tissue culture was performed as previously described (Risacher et al., 2009). Individual plantlets were transferred to Jiffy-7 pellets (LBS Horticulture) and acclimatised to ambient humidity, prior to potting into 9 cm pots containing M2 compost plus 5 g/L slow release fertilizer (Osmocote Exact 15:9:9). Plants were grown to maturity and seed harvest in controlled environment chambers, as above.

Quantitative PCR was used to determine T-DNA integration number based on an *nptII* copy number Taqman probe assay relative to a single copy wheat gene amplicon, GaMyb, normalised to a known single copy wheat line (Milner et al., 2018). Homozygous and null T1 plants were identified by nptII copy number Taqman probe assay as above with comparison to the predicted Mendelian inheritance ratios for 1- and 2-loci segregation, using a Chi squared test. Peroxisome fluorescence intensity and longevity at 18 days post-germination (the timing of *Z. tritici* infection) in subsidiary cells was only found to be sufficient in lines with more than 4 T-DNA insertions. Line CYT21.1 from this group was taken forward and was confirmed to have no obvious growth phenotype nor changes in compatibility towards the *Z. tritici* strains used in this study. This line contains two copies of the *nptII* gene at a single locus. T4 seed was used for all experiments.

### 2.4 Transmission light microscopy and analysis

Transmission microscopy experiments to investigate cell wall changes was performed on living wheat leaves infected with *Z. tritici* at 7 to 10 days post infection (DPI) during the asymptomatic phase. Stomatal responses to leaf infection were imaged using upright optical light transmission microscopy under 40x objective magnification and Koehler illumination with a wide condenser aperture. The leaves were mounted using perfluorocarbon PP11 (F2 Chemicals, UK) with the purpose of avoiding invasive clearing and staining procedures (Littlejohn et al., 2014). The images were taken using a colour CCD camera (Qimaging, Canada) and analysed using Fiji. Papilla-like body boundaries in subsidiary cells were defined manually using the polygon selection tool. The centroid (centre of mass) position and perimeter length were calculated from these regions of interest (ROIs). Positions along the aperture length of both hyphae and papillae were normalised to values between 0 and 1, with 0 and 1 representing the aperture ends. Datasets to simulate the null hypothesis of uncorrelated positions were generated using two methods: first by randomly generating paired locations between 0 and 1 and second by shuffling the measured hyphae and papillae pairs. The second method is limited by the complexity of each dataset but ensures only biologically valid positions are used. Normalised distances between hyphae and papillae were calculated and populations compared in MATLAB2021A using the Kruskal-Wallis test combined with *post-hoc* analysis that included Bonferroni corrections.

### 2.5 Histochemical and immunofluorescence imaging

Aniline blue was used to visualise callose deposition in infected wheat tissues. Inoculated leaves were cut into 2-cm pieces, mounted in the staining solution (0.1% w/v aniline blue in 150 mM K_2_HPO_4_ pH to 9.5) and vacuum infiltrated using a vacuum pump at room temperature. Wheat leaves infected with Z. *tritici* were collected 7 to 10 DPI and 16-40 hours after *Bgh* inoculation (HAI). Samples were viewed within 30 minutes using an inverted Leica TCS SP8 Confocal Laser Scanning Microscope with a 40x immersion oil lens and HyD detectors. Images were acquired using a 405 nm excitation laser combined with an emission window set to 430-500 nm (for aniline blue) and 670–756 nm (for chlorophyll autofluorescence). Stacks of optical sections were scanned at 700 Hz with a step size of 0.5 µm at 512×512 pixel resolution, using a pinhole of 1.0 Airy units.

Callose immunofluorescence was achieved using anti-callose monoclonal antibody 400-2 (Biosupplies, Australia) specific for (1→3)-β-glucans. Leaves infected with *Z. tritici* IPO323 were collected 9 DPI, fixed in PEM buffer (40 mM Pipes, 5mM EGTA, 5mM MgSO_4_ containing 4% (v/v) paraformaldehyde (Herve et al., 2010) and permeabilised by immersion in 0.5% v/v Triton X-100 in PBS (PBST) for 20 minutes. After washing three times in 1 x PBS the samples were incubated for 1 hour at room temperature with the primary antibody diluted 1:100 in PBS. After three washes with 1 x PBS the samples were incubated for 4 hours with anti-mouse secondary antibodies conjugated with Alexa Fluor 488 (ab150077, Abcam, UK) diluted 1:400 in PBS, then washed twice with PBST for 30 minutes. Samples were mounted using Fluoroshield histology mounting medium (Sigma-Aldrich, UK, Cat#F6182) and viewed under the microscope using a 488 nm laser and emission window 500-550 nm; 670-756 nm for chlorophyll autofluorescence.

### 2.6 Raman microscopy

Chemically specific imaging was performed using a Spectrally focused stimulated Raman scattering microscope based on the system described by Zeytunyan et al (Zeytunyan et al., 2018). The system is powered by a dual output fs laser (InSightX3 Newport spectra physics) which provides a pump beam at 802nm and a Stokes beam at 1045 nm, the beams are temporally and spatially overlapped and chirped to provide ps pulses in the SF-TRU spectral focusing unit (Newport spectra physics). This system also adds a 19.5 MHz modulation to the 1045nm beam required for SRS detection. This is then coupled to a modified confocal scanning microscope (Olympus FV3000) and the light is focused onto the sample with a 60x 1.2 NA objective (UPlanSApo 60x). SRS is detected in the forwards direction using a photodiode coupled to a lockIn amplifier (APE SRS detection set) with the following filters Chroma CARS 890-210 and Edmund optics 950 nm short pass filter) to exclude the 1045 nm Stokes beam. A series 101 of images is taken at Raman shifts ranging from 2790-3140 cm^-1^ to provide a hyperspectral image cube covering the CH vibrational region.

### 2.7 Live cell imaging of peroxisomes

Bobwhite S56 wheat lines expressing mCherry-SKL infected with either *Z. tritici* or *Bgh* were mounted using perfluorocarbon PP11 solution and viewed under an inverted Leica TCS SP8 Confocal Laser Scanning Microscope with a 40x immersion oil lens (excitation 561 nm, detection at 578-651 nm). Chlorophyll autofluorescence was detected in the range of 670–756 nm with a HyD detector using a 488 nm argon laser line. Three consecutive optical stacks were recorded at a scan speed of 700 Hz (resulting in approximately 60 s between each stack) with a step size of 0.64 µm at 256×256 pixel resolution, using a pinhole of 2.0 Airy units.

### 2.8 Peroxisome position analysis

Images were initially analysed using Fiji to identify the region of interest, which consisted of two guard cells and two subsidiary cells, named proximal or distal in relation to the hyphae direction of approach. Subsequent analysis was performed using MATLAB2016A (Mathworks, US). Full scripts can be downloaded as supplemental data (https://uk.mathworks.com/matlabcentral/fileexchange/114885-peroxisomestool). In brief a blob-finder algorithm was constructed to detect the presence of fluorescent spheroid organelles using a Laplacian of Gaussian (LoG) filter and tracked using an implementation of the Hungarian matching / Munkres algorithm. The algorithm tracked the identity of peroxisomes through each Z-stack. The spatiotemporal centroid of individual peroxisome tracks and the direction and the magnitude of peroxisome displacement was calculated and displayed using a custom MATLAB script. The average peroxisome position for each cell was defined as the mode of all peroxisome positions, which is more appropriate than the mean or median since the mean and median positions are sensitive to the relative peroxisome population size within the cluster, inaccurately displacing the cluster position towards the centre of the cell X-axis. Correlation between hyphal position Xhyp and modal peroxisome position Xper were tested by studying the distribution of |Xhyp-Xper|, the absolute value of the distance between the two positions. Whether this distribution showed any significance difference between genotypes or to a randomly-generated null distribution were tested using both the Kolmogorov–Smirnov and the Mann– Whitney U tests.

### 2.9 Plasmolysis of wheat epidermal cells

Bobwhite S56 mCherry-SKL leaves inoculated with *Z. tritici* IPO323 at 9 DPI were incubated in 5 M sorbitol (Sigma-Aldrich, UK, Cat#S1876) for one hour prior to mounting (in the same solution) and imaged using confocal microscopy.

## 3 Results

### 3.1 *Z. tritici* hyphae induce subsidiary cell papillae

We used light microscopy to test the hypothesis that wheat epidermal cells show positional sub-cellular responses to *Z. tritici* during surface hyphal growth and stomatal invasion. We used Cadenza, a cultivar known to carry an allele of the *Stb6* wall-associated kinase that confers a significant but imperfect level of resistance to IPO323 (Arraiano and Brown, 2006; Lee et al., 2015; Rudd et al., 2008). For comparison we used the cultivar Riband (with an insensitive *Stb6* haplotype) as an IPO323-compatible control. Additionally, we compared Bobwhite S56, a cultivar used by our laboratory for transgenic live-cell imaging. All cultivars compared against Riband showed resistance to IPO323 infection when quantifying scattered or clustered pycnidia on the adaxial surface of the infected leaves (Supplemental Figure 2A&B). Sequencing of PCR amplified fragments of the *Stb6* locus of Bobwhite S56 confirmed the absence of known *AvrStb6*-susceptible haplotypes (Saintenac et al., 2018). All of the wheat cultivars showed susceptibility following infection with *Z. tritici* field strain K4979N2X1 (henceforth abbreviated to N2X1; Supplementary Figure 2B).

We searched for microscopic early indicators of host defence responses in these cultivars using live-cell transmission light microscopy at 7-10 DPI, prior to the appearance of host symptoms indicative of the necrotrophic phase. We specifically applied measures to preserve spore and hyphal contact points by avoiding clearing and staining procedures and mounting samples directly using PP11, a low surface tension medium with a plant tissue-matched refractive index (Littlejohn et al., 2014). Most epidermal cell types of compatible and resistant cultivars did not show any transmission-detectable response to the presence of spores or hyphae. Cadenza and Bobwhite S56 cultivars formed pigmented, globular, papilla-like structures within the cytoplasm of cells within the stomatal cell complex (Figure 1A). This occurred in response to both avirulent IPO323 and virulent N2X1. These structures were distinct in morphology and cell specificity from those previously reported using clearing and staining procedures for fungal-induced cell wall modifications (Shetty et al., 2009, 2007, 2003). These bodies were most evident in subsidiary cells that flank the guard cells, irrespective of direct contact occurring between the observed penetrating hypha and subsidiary cell. To control for the potential bioactivity of PP11 in their formation we confirmed the presence of equivalent papillae structures in water-mounted samples (Supplemental Figure 2C). Analogous features were not identified in non-stomata epidermal cell types in contact with hyphae or spores in either of the mounting conditions. Less extensive non-pigmented papilla-like bodies were observed in the Riband cultivar when infected with either of the *Z. tritici* strains used in this study (Figure 1A). Approximately 25% of IPO323 interactions with Riband stomata resulted in papilla formation compared to approximately 60% of IPO323 in contact with Bobwhite S56 (n=16 and n=24 respectively) further suggesting that the host genotype can influence the manifestation of the subsidiary cell papilla response.

**Figure 1.**
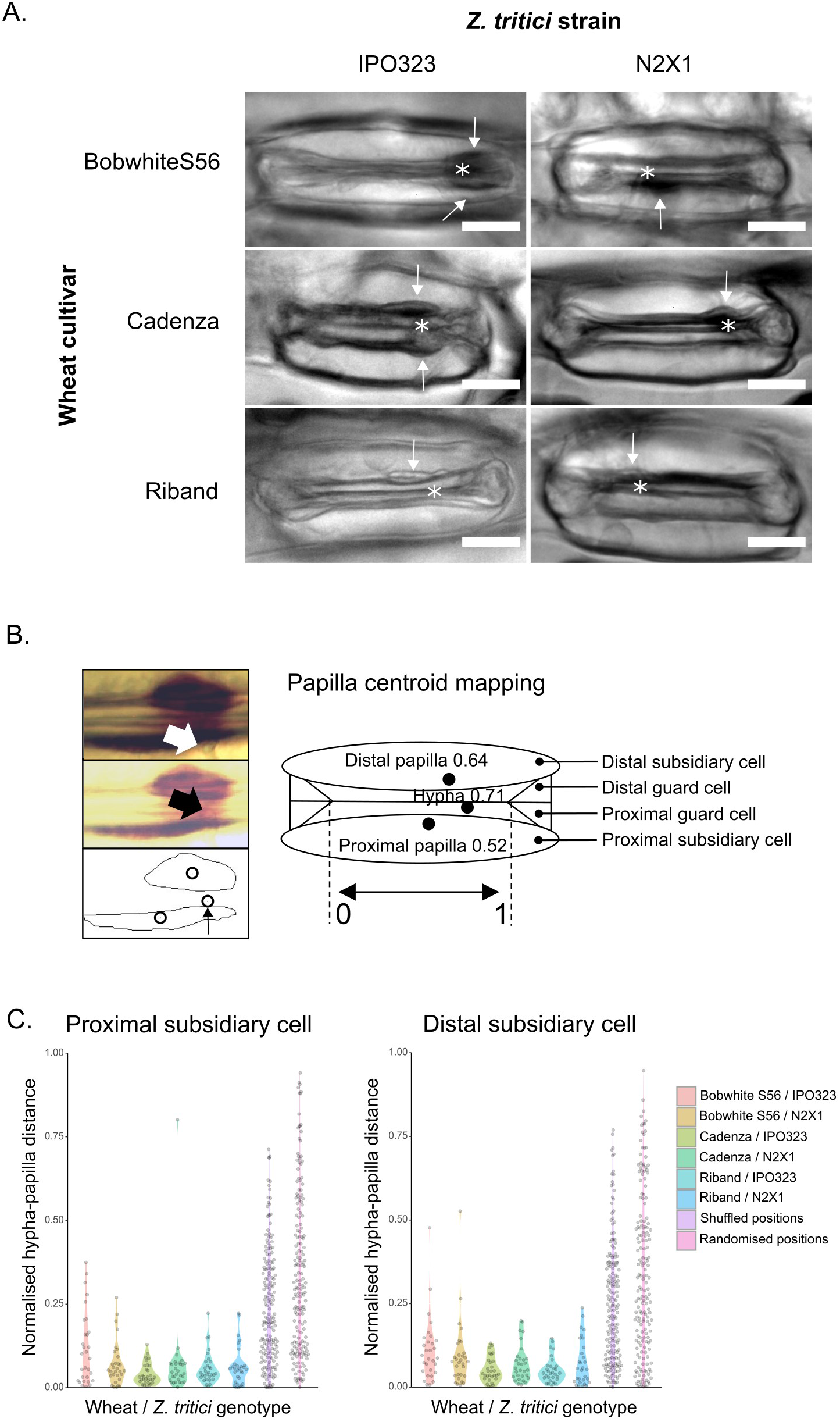
A. Papillae-like bodies (white arrows) formed at the subsidiary cells following hyphae detection at the guard cell entry (asterisks indicate hyphal transition). Bobwhite S56, Riband and Cadenza wheat leaves were challenged with *Z. tritici* IPO323 and N2X1. Stomatal responses to leaf infection were imaged 7-10 DPI. Scale bars are 20 μm. B. Schematic representation of the measurement of papillae-like body positions. Left panel shows papillae like-bodies and the hypha stimulus (white arrow). Lower left panel shows the centroid positions of the imaged papillae and the position of fungal ingress (also shown in the middle panel, indicated by black arrow). Right panel shows a representation of the labelling of the stomatal complex relative to fungal pre-invasion. Guard cells (GC) and subsidiary cells (SC) were named proximal and distal in relation to the hyphae direction of approach. Coordinates along the stomatal complex X-axis were normalised between 0 and 1. C. Plot showing the distance calculated between the papillae centroid X positions and hyphal X positions in cultivars Bobwhite S56, Riband and Cadenza combined with two *Z. tritici* strains IPO323 and N2X1 in both proximal and distal subsidiary cells. ‘Shuffled positions’ were calculated by randomly pairing hyphal and papilla positions, while ‘randomised positions’ show distances generated from random placement of hyphae and papillae in positions between 0 and 1 (please see materials and methods for details).

### 3.2 Subsidiary cell papillae are positioned with precision beneath fungal hyphae

We then asked whether the positions of papilla-like bodies were dependent upon the precise location of the hypha at the aperture. In order to describe the identity of different cells within the stomatal complex during immune responses to *Z. tritici*, we established a labelling-scheme based upon the direction of approach of each invading fungal hypha. The wheat guard cell complex consists of two guard cells and two subsidiary cells that we designated as either proximal or distal depending on the hyphal direction of approach. Papilla locations were mapped by manually tracing the boundaries of pigmented bodies and calculating a centroid position (Figure 1B). The co-ordinates were normalised between stomata by defining the left-most boundary of the aperture with a positional value of ‘0’ and the right-most with a value of ‘1’. Papilla centroid positions showed strong positive correlations between the relative X position of both proximal and distal papillae centroids and the X position of hyphae transiting through the stomatal aperture. This was true for all six host-microbe combinations, with Pearson correlation values ranging from 0.71 to 0.98 (supplemental figure 2D). We calculated the normalised distances between hyphae and papillae and compared these values to randomised and shuffled datasets to simulate the null hypothesis (an absence of correlation). All genotype combinations were significantly different from both null comparators (Figure 1C; p< 0.005). No significant difference was detected between genotype combinations suggesting that papilla position was not influenced by host or pathogen genotype.

### 3.3 Callose encapsulates distinct papilla content

We used the callose stain aniline blue on Bobwhite S56 wheat leaves infected with *Z. tritici* IPO323 to further investigate the molecular composition of the papilla like-bodies. We also imaged the responses to *Bgh* appressoria as a positive control, as focal responses to this fungus are relatively well understood and known to contain callose (Chowdhury et al., 2014) but to our knowledge such responses in subsidiary cells have not been described previously. We found that *Bgh* stimulated papilla formation in subsidiary cells and these were enriched with callose (Figure 2A). The papilla-like bodies produced in response to IPO323 showed aniline blue fluorescence specifically at the papilla boundary (Figure 2B), in a capsule-like formation distinct from papillae formed in response to *Bgh*. Furthermore, IPO323-stimulated papillae did not show a strong autofluorescence signal suggesting distinct contents compared to *Bgh*-induced papillae. We then used immunofluorescence to confirm the boundary composition of the papilla-like-bodies. Consistent with aniline blue fluorescence, the anti-callose antibody was detected specifically at the periphery (Figure 2C).

**Figure 2.**
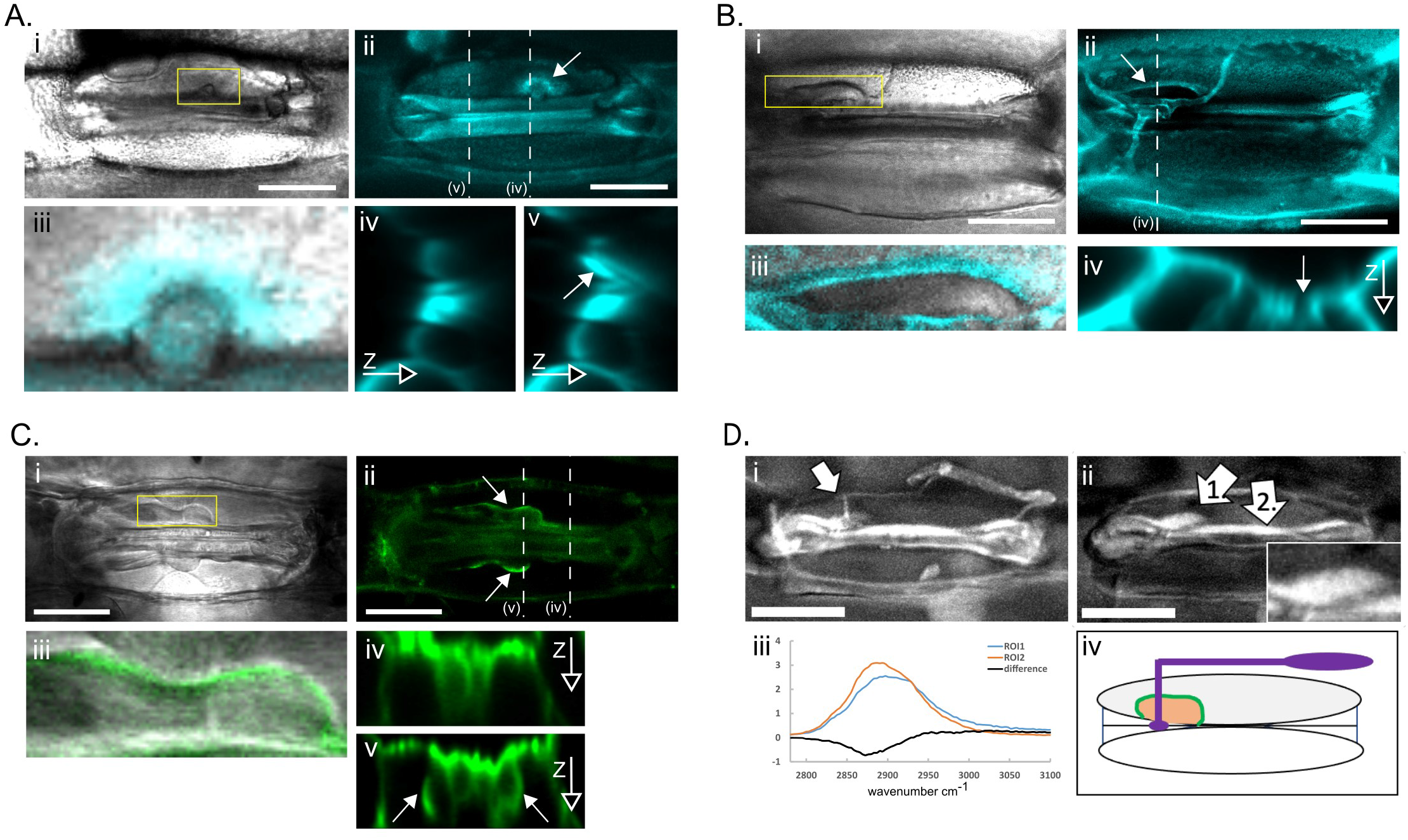
A. Callose labelling with aniline blue of Bobwhite S56 wheat subsidiary cells interacting with *Blumeria graminis f. sp. hordei* 24 h after inoculation (HAI). (i) Transmission image of wheat stoma showing papilla formation associated with a fungal appressorium (boxed area). (ii) Panel showing fluorescence signal with arrow indicating callose deposition. (iii) Composite image showing zoomed papilla-like body and fluorescence. (iv) Transverse optical projections of the same guard cell complex along the dotted line indicated in panel ii. (v) Comparable optical projection through the papilla (indicated by white arrow). B. Callose labelling of Bobwhite S56 wheat leaves challenged with *Z. tritici* IPO323 hypha 9 days post infection. (i) Transmission image of wheat stoma showing a papilla-like body (boxed area) adjacent to penetrating hypha. (ii) Panel showing fluorescence signal (white arrow indicates callose deposition at the periphery). (iii) Composite image showing zoomed papilla-like body with callose periphery. (iv) Transverse optical projection through the callose deposit showing the unstained papilla interior (indicated by white arrow). C. Indirect immunofluorescence for callose using Bobwhite S56 wheat leaves challenged with *Z. tritici* IPO323 9 days post infection. (i) Transmission image of wheat stoma showing papilla-like bodies (boxed area indicates a single papilla). (ii) Green fluorescence channel showing secondary antibody signal (white arrows indicate papillae periphery). (iii) Composite image of boxed area. (iv) Transverse optical projections of the same guard cell complex along the dotted line indicated in panel ii. (v) Comparable optical projection through the papillae (indicated by white arrows). D. Label-free CRS imaging in the CH band highlights the invading fungal network in wheat tissues (i) and CH-rich papilla content within the callose capsule (ii; arrow ‘1’ and inset). The CRS spectra (iii) show distinct profiles when comparing papilla content to neighbouring cell walls (‘1’ vs ‘2’). The combined information from the different probe and imaging techniques is summarised in the scheme in iv (purple = fungal network, green =callose encapsulated papilla, orange = CH-rich papilla content). All scale bars are 20 μm.

Our histological and immunofluorescence approaches failed to label the content of the callose capsule and no autofluorescence was detected above cytoplasmic background in any band using laser-scanning confocal microscopy. Therefore, we used label-free coherent Raman scattering (CRS) microscopy to compare papilla content with the cytoplasm and neighbouring cell walls. CRS imaging in the CH band (from 2800-3100 cm^-1^) highlighted both fungal and host structures (Figure 2D). The signal from host subsidiary cells was concentrated to the cell walls and papilla with cytoplasmic signal largely restricted to the nuclei. The CRS spectra showed distinct profiles when comparing papilla content to neighbouring cell walls suggesting that subsidiary cell papillae induced by *Z. tritici* have a metabolite profile distinct from the cytoplasm and other neighbouring cell wall content.

### 3.4 mCherry-SKL labels peroxisomes within wheat subsidiary cells

In order to test the hypothesis that cell wall modification was accompanied and/or preceded by changes in subcellular dynamics, we developed stable transgenic wheat lines expressing peroxisome-targeted fusion protein mCherry-SKL (Nelson et al., 2007) under the OsActin promoter (Supplementary Figure 1&3). We used mCherry-SKL labelled peroxisomes to further test the hypothesis that the papilla-like bodies in subsidiary cells are associated with the cell wall rather than the cytoplasm. Peroxisomes were not observed to cross the papilla boundary. Moreover, we induced plasmolysis using 5 M sorbitol solution in wheat leaves infected with the *Z. tritici* and observed that papillae remained attached to cell walls rather than shrinking with the peroxisome-labelled cytoplasmic compartment (Supplementary Figure 4). The suitability of our transgenic line for testing the presence of pathogen-induced organelle distributions in subsidiary cells was validated by performing confocal microscopy on wheat leaves infected with *Bgh*. Powdery mildews are known to induce pronounced perturbations in peroxisome and other organelle distributions in both dicots and monocots (An et al., 2006; Koh et al., 2005), but this has not been previously tested for in subsidiary cells. Peroxisome distribution was imaged by collecting Z-series (three consecutive stacks) of optical sections through stomatal complexes interacting with *Bgh* appressoria (Figure 3). Our analysis identified a correlation between the modal peroxisome position and appressorium position along the X-axis (n = 10 cells; Pearson’s correlation coefficient of 0.88). This showed that subsidiary cells respond to *Bgh* by diverting a population of peroxisomes to the site of interaction and therefore could have the capacity to respond to *Z. tritici* in an equivalent manner.

**Figure 3.**
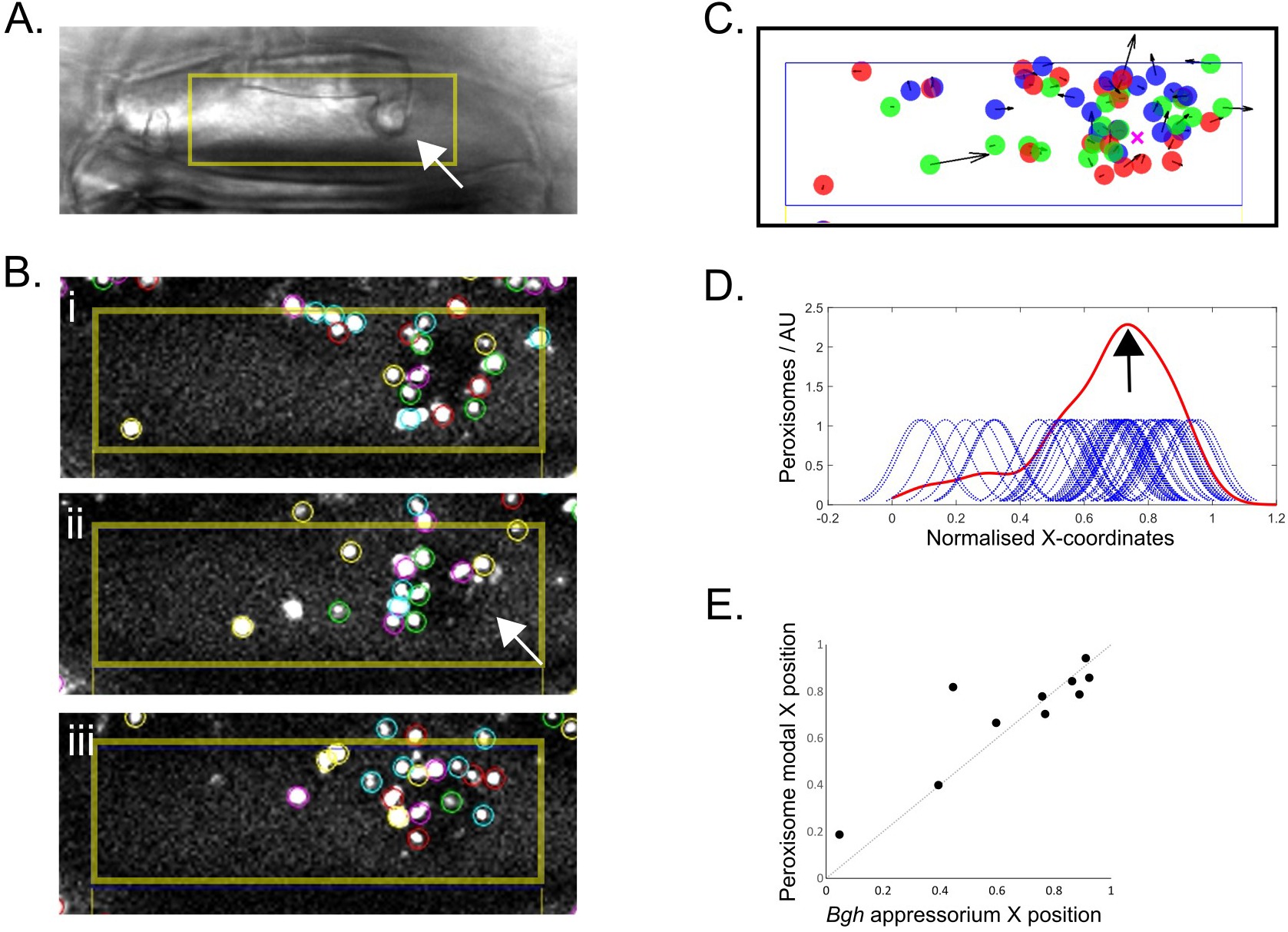
A. Transmission channel image showing a *Bgh*-inoculated subsidiary cell. White arrow indicates the position of the fungal appressorium and host papilla. The yellow box denotes the boundaries of the area used in the subsequent analysis. B. Fluorescent objects were identified, tracked during Z-stack progression and filtered by track-length to ensure high-stringency detection of peroxisomes. Three Z-stacks (single optical slices of which are shown in panels i, ii and iii) were obtained per sample. Coloured circles indicate algorithm-detected peroxisomes. C. Plot of detected peroxisomes in B. The blue box is equivalent to the yellow box in A and B. Coloured dots represent the mean position of each tracked peroxisome with red, green and blue colours referring to peroxisomes tracked in Z-stack 1, 2 and 3 respectively. The pink cross indicates the appressorium position. D. Gaussian Kernels representing each peroxisome on the X-axis were plotted to produce a smooth histogram. The modal peroxisome position is indicated with a black arrow. E. Scatter plot showing the relationship between normalised appressorium X co-ordinates and the peroxisome modal X position (n=10; Pearson’s correlation coefficient = 0.88). X positions within the subsidiary cell areas were normalised between 0 and 1.

### 3.5 Peroxisome distribution is influenced by hyphal position

Peroxisome-labelled wheat leaves were infected with *Z. tritici* strains IPO323 and N2X1 and were viewed using confocal microscopy at 3 to 5 DPI, with the aim of imaging hyphae and host cells at a timepoint close to the event of aperture-transit. A minimum of 15 transit sites were analysed for proximal and distal cell positions combined with either IPO323 or N2X1 hyphae. Peroxisome distribution and its correlation with hyphal position appeared to vary considerably on a cell-by-cell basis with a spectrum of association ranging from distinct tight clusters at hyphal ingression sites (Figure 4A) to unaligned clusters and wider distribution of peroxisomes (Figure 4B). We measured the correlation between the modal position of the peroxisome population and the position of the hypha along the X-axis of the stomatal aperture. When pooling data for both genotypes and cell types a weak Pearson correlation of 0.44 between modal position and hypha position was evident (Figure 4C). We tested the significance of this association by calculating the normalised measured distances between hyphae and modal positions and comparing these to the distances that could be calculated from random pairings of positions (Figure 4D). Both a Mann-Whitney U test and a Kolmogorov-Smirnov test show significant differences between the measured population and simulated null distances (p = 4.9 × 10^−3^ and p = 8.3 × 10^−3^ respectively), supporting a correlation between positions. A three-way comparison between IPO323, N2X1 and the shuffled dataset (using the Kruskal-Wallis test) revealed no significant differences, suggesting that the genotype of the hypha does not have a strong impact upon the levels of peroxisome-hypha association.

**Figure 4.**
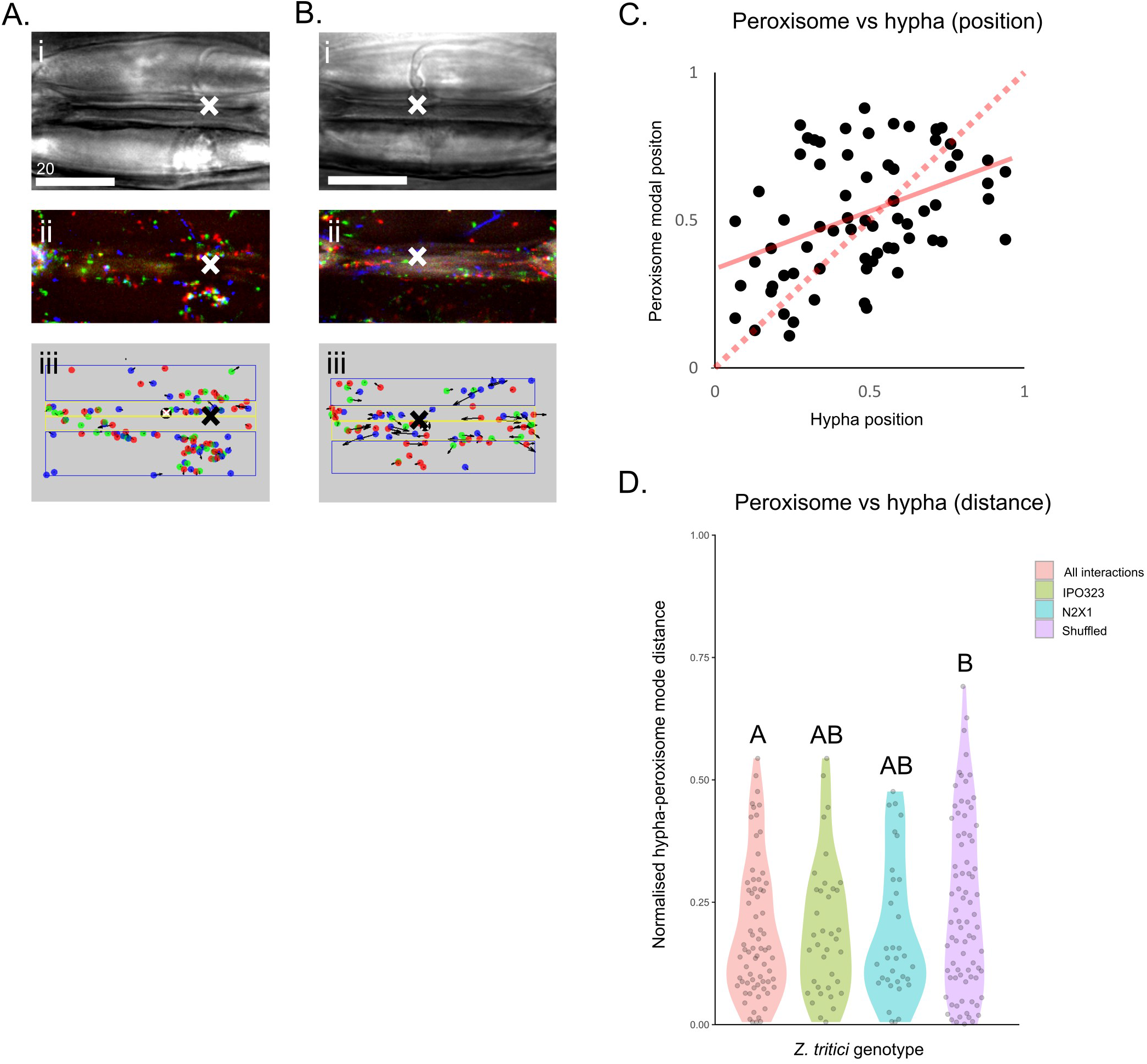
A&B. Confocal images of Bobwhite S56 wheat stomata inoculated with *Z. tritici* with examples of subsidiary cell peroxisomes clustering at the site of hyphal contact (A) and absence of pronounced hypha-correlated peroxisome clustering (B). Panels i show transmission images, panels ii show maximum projections of peroxisome populations with colour coding showing data from pass 1 (red), pass 2 (green) and pass 3 (blue). Panels iii show the extracted peroxisome positions that correspond with the source data shown in ii. Blue boxes show the quantified areas of the subsidiary cells, yellow boxes represent the guard cells. Coloured dots represent the mean position of each peroxisome track. White and black crosses show the ingression site of the relevant hypha. Scale bars are 20 μm. C. Scatter plot of normalised X *Z. tritici* hyphae position vs peroxisome modal X position including both IPO323 and N2X1 data (n=80). These data show a Pearson correlation value of 0.44 with the dashed red line representing a Pearson correlation of 1 for comparison. The solid red line shows a linear regression calculated from the data with an R^2^ value of 0.19. D. Violin jitter plot showing the distance calculated between peroxisome X positions and hyphal X positions. Pooled data between the two strains shows a statistically significant shortening of the distances between measured hypha and peroxisome mode pairings compared to shuffled locations. Individual strain-specific datasets do not show statistical difference from the control or between each other.

## 4. Discussion

Host cell wall changes at the site of invasive microbial contact have been observed across the plant kingdom and in response to a broad range of microbial life including bacteria, oomycetes and fungi. The focusing of cytoplasmic resources and targeted reinforcement of cell walls are hallmarks of these responses and are considered to enable repulsion of a potential pathogen through a mixture of mechanical and chemical defence. For example, the ABC transporter protein PEN3 of *Arabidopsis thaliana* is enriched at microbial contacts and has been shown to enable the release of fungicidal secondary metabolites that make a significant contribution to non-host responses towards barley-adapted powdery mildew and other microbes (Lu et al., 2015). PEN3 function also contributes to callose deposition to alter the local physical properties of the plant cell wall (Matern et al., 2019). Enhancing and reinforcing these mechanisms could provide new approaches to crop protection to focus these responses towards virulent microbes that typically circumvent the immune systems of their natural hosts.

The fungal phytopathogen *Z. tritici* evades the wheat immune system for long periods of time following its ingress into leaf air spaces and shows the ability to colonise large volumes of host tissue with minimal detectable response from the plant (Marshall et al., 2011). Here we have used minimally invasive light microscopy methods to show that despite this ‘stealth’ behaviour *Z. tritici* can stimulate some features of focal immune responses within compatible and incompatible hosts. Neither blastospore germination nor hyphal growth across the leaf surface were observed to stimulate epidermal cell wall reinforcement or the accumulation of cytoplasmic masses typically observed during focal immune responses. However, transgression of the stomatal aperture by *Z. tritici* hyphae was seen to trigger spatially-focused dynamics in host cells. Subsidiary cells flanking guard cells developed cell wall papillae spatially correlated with the point of ingress in multiple host-pathogen genotype combinations. These papillae show atypical characteristics when compared to papillae generated by the same cell type in response to powdery mildew. Callose deposition was restricted to the surface layer of the papillae with the main bulk consisting of unknown content with minimal levels of autofluorescence. By contrast, papillae associated with powdery mildew were enriched at their core with callose and showed prominent autofluorescence, typical of papillae grown in response to cell-penetrating hyphae (Chowdhury et al., 2014). Subsidiary cell papillae are therefore products strongly shaped by the exact identity or the invasion-style of the interacting hypha.

Wheat shows gene-for-gene resistance behaviour towards *Z. tritici. Stb6* was the first of these genes to be cloned and was found to encode a cell wall associated kinase that interacts genetically with the apoplastic effector *AvrStb6* (Saintenac et al., 2018). Several members of the plant WAK family interact with host cell wall-derived carbohydrates and are thought to report on the integrity of the plant cell wall (Kohorn, 2016; Saintenac et al., 2021). Recently a second wheat resistance gene, *Stb16*, was identified as a cysteine-rich receptor like kinase (CRK) that also potentially binds carbohydrates to trigger resistance to specific strains of *Z. tritici* (Saintenac et al., 2021). Both *Stb6* and *Stb16* arrest growth at relatively early stages of infection and encode plasma membrane-localised kinases with extracellular receptor domains (Saintenac et al., 2021, 2018). Whatever the identities of their unknown ligands, these receptor-encoding resistance genes suggest that extracellular awareness of *Z. tritici* is critical for successful defence. Our data indicate that the perception of *Z. tritici* extends to a sub-cellular resolution, as both papillae and organelle distribution in host subsidiary cells track hyphal location. This, however, is to some extent independent of the gene-for-gene interactions as our observations show that both susceptible and resistant haplotypes of *Stb6* generate hypha-associated papillae when combined with either compatible or incompatible strains of *Z. tritici*. Moreover, organelle redistribution does not differ significantly between compatible and incompatible stimuli. These responses therefore reveal the capacity of host cells to locate hyphal stimuli. In the compatible interactions we have observed that *Z. tritici* hyphae are able to progress despite this response. This suggests that in isolation this response is not sufficient to prevent disease.

### 4.1 Subsidiary cells are associated with immune responses

Stomata in wheat consist of a pair of subsidiary cells flanking two guard cells that coordinate osmotic pressure changes to regulate the aperture of stomatal pores (Franks and Farquhar, 2007; Mumm et al., 2011). The discovery of subsidiary cell wall modifications parallel to the location of hyphal ingress through the pore implies a secondary function for subsidiary cells in mediating plant-microbe interactions. The responses we have observed could be a combined product of host defence and pathogen virulence mechanisms, potentially obscuring their function, but their presence regardless suggests that boundaries of subsidiary cells are sensitive to fungal interactions. This highlights a knowledge-gap in the understanding of subsidiary cell identity and relevance to crop protection. Seminal contributions have been made by Raissig and colleagues towards understanding the role of the subsidiary cells through the use of the model grass *Brachypodium distachyon* (Raissig et al., 2017). A forward genetics screen for genes affecting stomatal morphology identified a subsidiary cells defective (*sid*) mutant unable to differentiate subsidiary cells. Stomatal responsiveness to different light conditions was significantly reduced in the mutant, suggesting that the absence of this cell type compromised the regulation of stomatal aperture (Raissig et al., 2017). Future exploitation of this mutant could also help reveal how subsidiary cells contribute to defence against microbes.

We labelled subsidiary cell papillae as either ‘proximal’ or ‘distal’ depending on the direction of hypha approach. We were surprised that, despite having no direct contact with the approaching hypha, distal cells were as accurate in the placement of their papillae as proximal cells. A complete guard cell interrupts the line of contact between hypha and proximal subsidiary cell. Theoretically hyphal position could be flagged by molecular pattern diffusion, by a secondary signalling chain across the guard cell, by pathogen-induced enzymatic cell wall irritation or by physical distortion of the subsidiary cell boundary by the clamped hypha. Overcoming the challenges of forward genetics screens in bread wheat, or future utilisation of the *B. distachyon* – *Z. tritici* non-host model are possible routes to discovering the mechanism behind this detection-at-a-distance phenomenon.

### 4.2 Peroxisome distribution as evidence for focal immune responses during wheat-*Zymoseptoria* interactions

Chloroplasts and nuclei of mesophyll cells have previously been reported to show altered morphologies during *Z. tritici* interactions (Kema et al., 1996). However, to our knowledge, no previous research has investigated changes in organelle distributions at the epidermis in response to *Z. tritici* invasive hyphae. Our observations of cell wall modifications in subsidiary cells required new analysis methods to test the hypothesis that these responses were concurrent with dynamic changes in organelle distribution, another hallmark of focal immune responses to fungi (Lipka et al., 2010). We developed a novel strategy for four-dimensional tracking (volume and time) of wheat peroxisomes labelled with mCherry-SKL (Nelson et al., 2007). We first validated the methodology by measuring peroxisome distribution within subsidiary cells responding to *Bgh* appressoria; a well-characterised stimulus for focal immune responses. These data confirmed peroxisome clustering around *Bgh* appressoria within subsidiary cells. We then applied this method to quantify organelle distribution during responses to virulent and avirulent strains of *Z. tritici*. However, no significant difference in the response based upon virulence could be identified. We used a stringent non-parametric comparison of measured hyphal-peroxisome paired co-ordinates against shuffled partnerships. However, it was clear both from visual inspection and detailed analysis that stomatal cells varied considerably in the levels of peroxisome clustering around hyphal sites. This could be due to a variation from site-to-site in the ‘awareness’ of hyphal presence or a consequence of a narrow response window. Such clustering was not observed in older 10 to 11 DPI samples where papillae were manifest, possibly supporting the concept of a finite window of organelle response.

Plant organelle movement and positioning are associated with external stimuli, including phytopathogens (Perico and Sparkes, 2018). Peroxisomes are key organelles that play vital roles in metabolism, ROS detoxification and signalling, and have been previously shown to aggregate at penetration sites of *Erysiphe cichoracearum* (Koh et al., 2005). In addition, multiple recent studies have shown the targeting of effectors by *Colletotrichum higginsianum, Magnaporthe oryzae* and powdery mildew to peroxisomes (Ning et al., 2022; Robin et al., 2018). Greater numbers of peroxisomes per cell may be associated with successful defence of wheat against Wheat Streak Mosaic Virus (Mishchenko et al., 2021) possibly revealing a defence trait that can be linked to peroxisome behaviour. The physiological functions of peroxisomes are diverse, with several directly impacting plant-microbe interactions. Peroxisomes are a site of jasmonic acid (JA) production (Theodoulou et al., 2005) and an absence of peroxisome catalase activity can trigger salicyclic acid (SA) production (Chaouch et al., 2010) suggesting that hydrogen peroxide dynamics mediated by peroxisomes are a significant factor in promoting systemic acquired resistance. The use of peroxisomes as a site of antimicrobial production has been proposed with the penetration-defence glycosyl hydrolase PEN2 observed to localise to peroxisomes (Lipka et al., 2008). More recently the functionally dominant localisation of PEN2 has been identified as being a mitochondrial rather than peroxisomal pool (Fuchs et al., 2016). This does not preclude the possibility that other phytoalexins and defensive metabolites could depend upon peroxisome-localised synthesis. The emergence of a local papilla could reflect two independent responses or the co-ordinated creation of an apoplastic reservoir for fungicidal products.

In conclusion, the ‘stealth’ pre-symptomatic pathogenesis of *Z. tritici* has the capacity to be perceived by the wheat immune system, to the extent that sub-cellular dynamics and cell wall growth respond locally to the fungal stimulus. These responses in themselves are not effective in preventing disease and may be, to some extent, a product of pathogen virulence. However these actions by the host could provide a kernel for engineering enhanced defence using the exemplar of *mlo*-enhanced penetration resistance in cereals as a template.

## Acknowledgements

This work was funded by BBSRC grants BB/H017569/1, BB/H018379/1 and BB/ M024172/1. MJD and DMR were supported by a Wellcome Trust Institutional Strategic Support Award (WT105618MA). DMR gratefully acknowledges financial support from the Medical Research Council (MR/P022405/1). We would like to thank Rhian Howells (NIAB), Graham Thomas (UoE), Harry Osborne (UoE) and Yaadwinder Sidhu (UoE) for training and technical assistance, Mike Csukai (Syngenta) for *Z. tritici* strain K4979N2X1 and Helen Fones (UoE) for comments on the manuscript.

## Author contributions

MJD, FV, AG, EW, JMo, KH and FV designed the research; FV, JMa, MJD, CR, EW, MC and SB performed the laboratory research; MJD, DMR, JMe designed the data analysis; GR and DH programmed the data analysis; FV and MJD wrote the manuscript with contributions from EW, DMR and JMa.

## Figure Legends

**Supplemental Figure 1**

Diagram showing the T-DNA regions of plasmids pEW235 and pEW242.

**Supplemental Figure 2**

A. Wheat leaves 24 days post *Z. tritici* IPO323 inoculation. Riband (i), Cadenza (ii), Bobwhite S56 (iii) and Riband mock-treated (iv). B. Bar charts showing percentage of inoculated wheat leaf area occupied by IPO323 or N2X1 lesions at 24 DPI. Zones of leaf tissue were classified as having healthy appearance, lesions with pycnidia or damage of indeterminate cause. Error bars represent the standard error from 3 biological repeats each containing a minimum of three inoculated plants of each combination. C. Bobwhite S56 wheat stomata with *Z. tritici* IPO323 transiting hypha showing proximal and distal papilla-like body (white arrows) mounted in water. Scale bar 20 µm. D. Scatter plots i to vi show the proximal papillae-like bodies normalised X positions (ordinate axis) vs. the hyphae normalised X positions (abscissa axis) in Bobwhite S56 (i, iv), Riband (ii, v) and Cadenza (iii, vi) wheat cultivars in response to *Z. tritici* IPO323 (i, ii, iii) and N2X1 (iv, v, vi) strains. Scatter plots vii to xii show the equivalent data for distal papillae-like bodies in Bobwhite S56 (vii, x), Riband (viii, xi) and Cadenza (ix, xii) wheat cultivars in response to *Z. tritici* IPO323 (xii, xiii, ix) and N2X1 (x, xi, xii) strains. Pearson values and p-values are highlighted in red and black respectively.

**Supplemental Figure 3**.

Labelling of F-actin and peroxisomes in transiently transformed wheat epidermal cells shows F-actin-associated peroxisome dynamics. A. Single confocal frame from a time series of an 84 × 17 μm cortical area of an epidermal cell co-expressing GFP-Lifeact and mCherry-SKL. Multiple time points within the boxed area are shown in the next panel. B. mCherry-SKL-labelled peroxisomes associate with actin GFP-Lifeact-labelled filaments while in motion. C, Mean projection of 100 frames (encompassing 152 seconds) of the GFP-Lifeact channel with superimposed movement vectors of the peroxisome population (contiguous tracks of length 4 or greater) within the same time period. White arrows indicate peroxisome points of entry into the field of view. D. Histogram of distribution of measured peroxisome speeds within a single cell (10 peroxisomes for 10 time intervals of 0.55 seconds). Scale bar is 20 μm.

**Supplementary Figure 4**.

Bobwhite S56 mCherry-SKL wheat stomata inoculated with *Z. tritici* IPO323 were treated at 9DPI for one-hour with 5 M sorbitol to induce plasmolysis. A. Brightfield image showing untreated stomatal cells with a subsidiary cell papilla (black arrow) formed in response to a *Z. tritici* hypha. B. Fluorescence channel image of the same cells showing peroxisomes labelled with mCherry-SKL (a single isolated peroxisome is indicated with a white circle). C. Brightfield image showing stomatal cells from the same leaf 1 hour post 5 M sorbitol treatment. The region of interest marked with a yellow square contains a cell wall papilla (black arrow) and plasmolysed protoplast (white arrow). D. Expanded brightfield image of the region of interest in C. The papilla remains attached to the subsidiary cell wall while the protoplast has retracted from the cell walls and papilla boundary. E. Fluorescence channel image showing peroxisomes within the boundary of the shrunken protoplast and their exclusion from the papilla. F. Composite image of D and E. Scale bars are 20 μm.

## Notes

https://uk.mathworks.com/matlabcentral/fileexchange/114885-peroxisomestool

### Competing Interest Statement

The authors have declared no competing interest.

