## Supplemental figures and table for "Wheat cells show positional responses to invasive *Zymoseptoria tritici*"

### Supplemental Figure 1

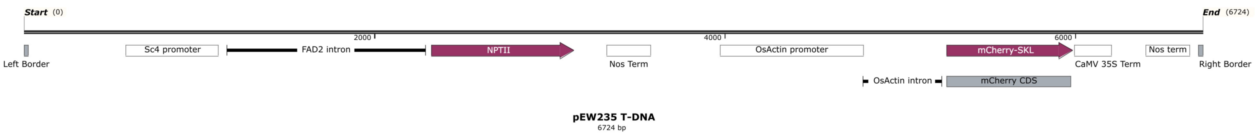


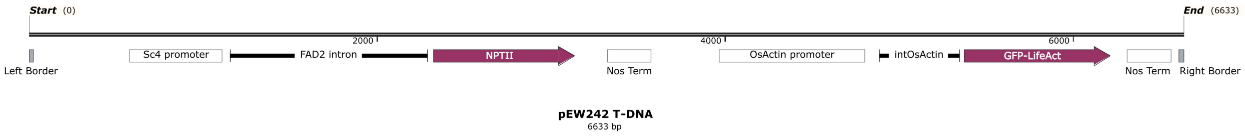

### Supplemental Figure 2

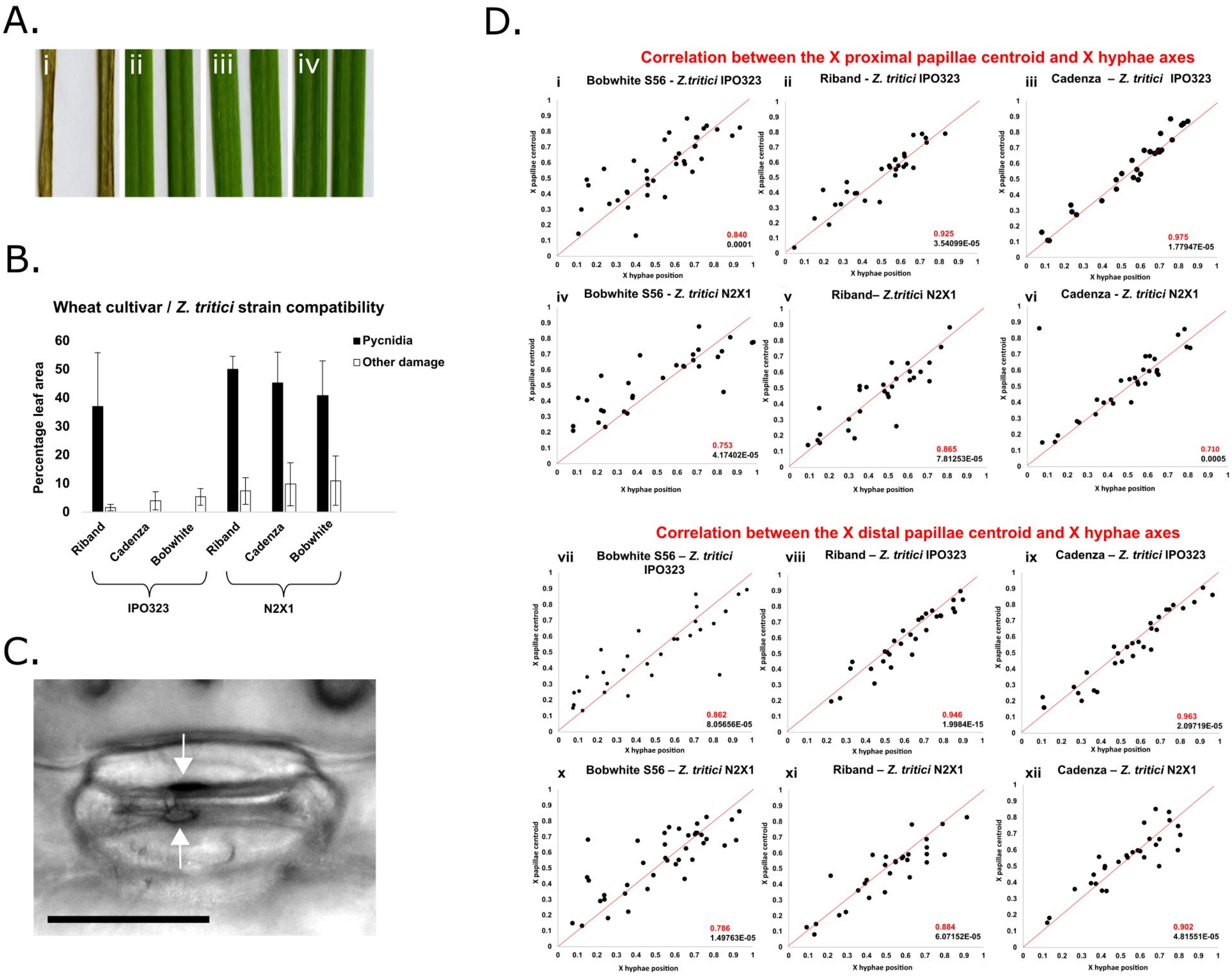

### Supplemental Figure 3

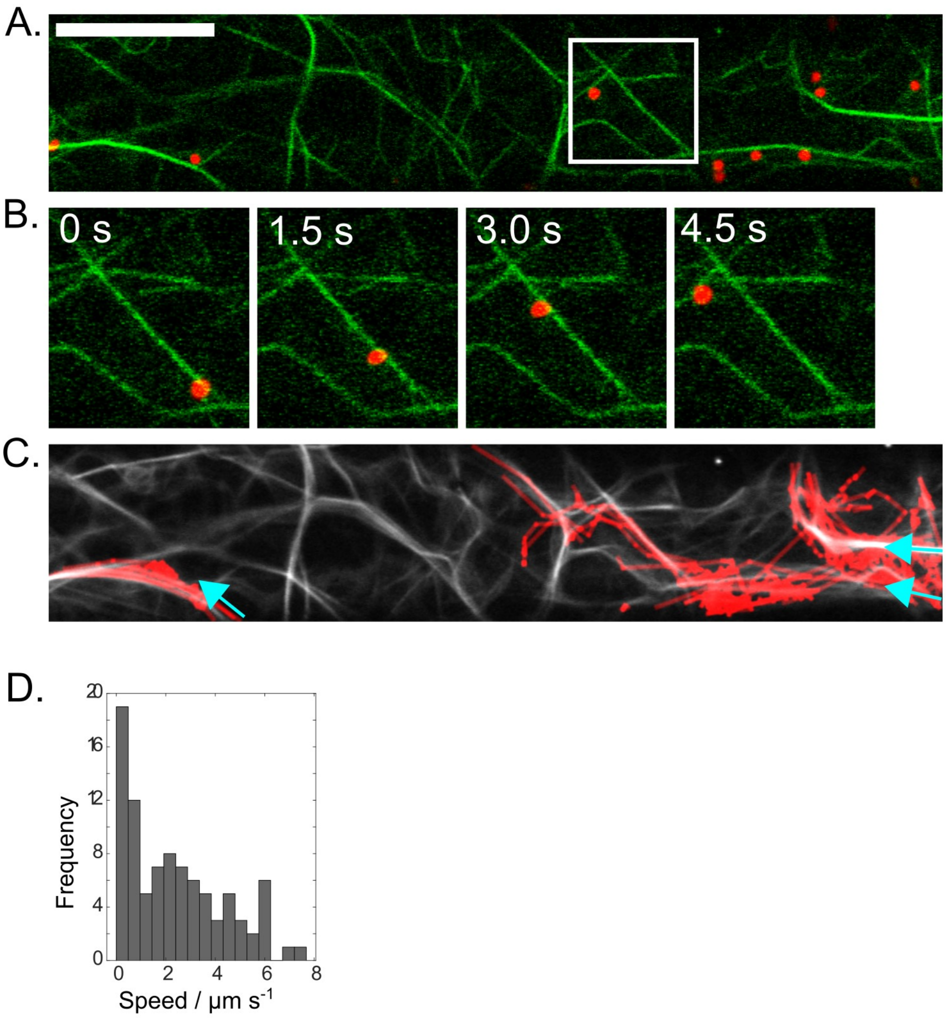

### Supplemental Figure 4

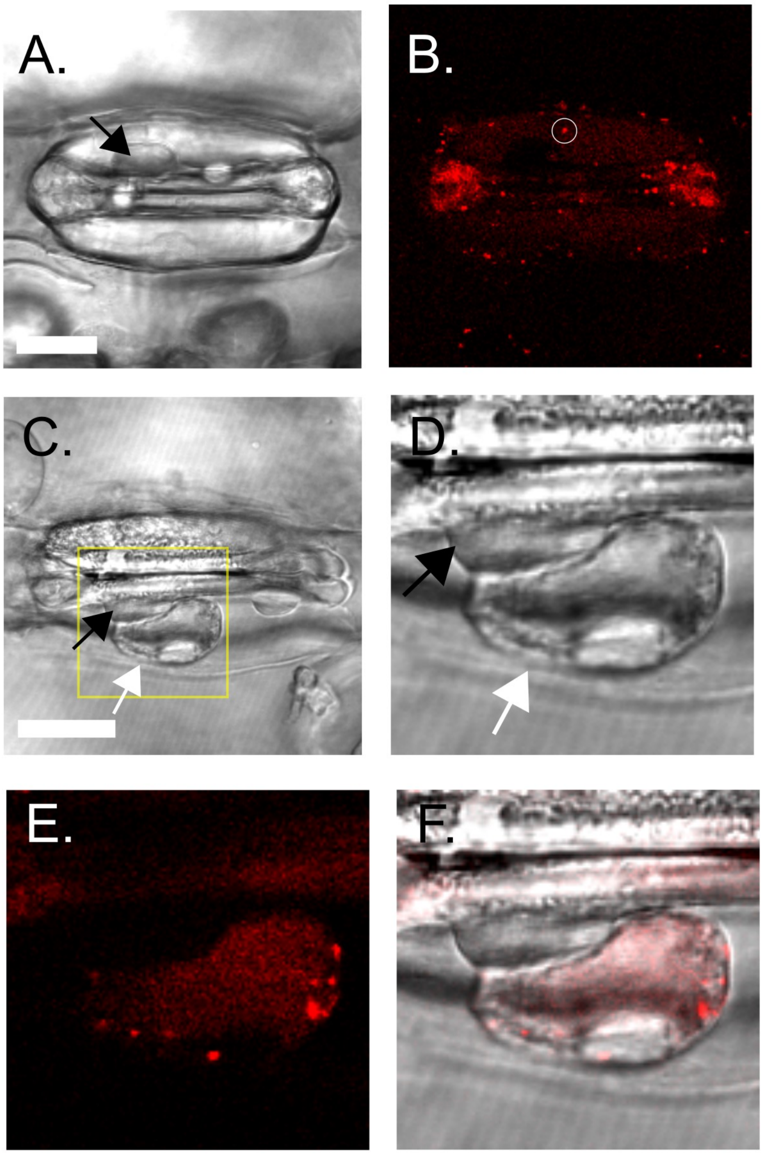

**Supplemental Table 1**

| **Name** | **Purpose** | **Sequence 5'-3'** |
|---|---|---|
| MDC43-for | pEW242 cloning | CCACCATGGTTAAAGGAGAAGAACTTTTCACTGG |
| MDC43-rev | pEW242 cloning | AGACTCGAGGGCGCGCCTTTGTATAGTTCAT |
| F2_Stb6 | Stb6 genotyping | GATGGTCGTCTGGTTGCAGT |
| R2_Stb6 | Stb6 genotyping | ATACCCTTGCATCGTCCTACT |
| F4_Stb6 | Stb6 genotyping | AAAGGTGGTTACGGTGTGGT |
| R4_Stb6 | Stb6 genotyping | AAGATCGGAGGAGACAAGCC |
| F2_Stb6kin | Stb6 genotyping | CCTTCCGTGTTCCCACTGTTG |
| R2_Stb6kin | Stb6 genotyping | AGGAGCTAGAGGAACAATTG |

PCR and sequencing primers used in this study.
